# Millisecond-scale molecular dynamics simulation of spike RBD structure reveals evolutionary adaption of SARS-CoV-2 to stably bind ACE2

**DOI:** 10.1101/2020.12.11.422055

**Authors:** Gard Nelson, Oleksandr Buzko, Aaron Bassett, Patricia Spilman, Kayvan Niazi, Shahrooz Rabizadeh, Patrick Soon-Shiong

## Abstract

The Receptor Binding Domain (RBD) of the SARS-CoV-2 surface spike (S) protein interacts with host angiotensin converting enzyme 2 (ACE2) to gain entry to host cells and initiate infection^1–3^. Detailed, accurate understanding of key interactions between S RBD and ACE2 provides critical information that may be leveraged in the development of strategies for the prevention and treatment of COVID-19. Utilizing the published sequences and cryo-EM structures of both the viral S RBD and ACE2^4,5^, we performed *in silico* molecular dynamics (MD) simulations of free S RBD and of its interaction with ACE2 over the exceptionally long durations of 2.9 and 2 milliseconds, respectively, to elucidate the nature and relative affinity of S RBD surface residues for the ACE2 binding region. Our findings reveal that free S RBD has assumed an optimized ACE2 binding-ready conformation, incurring little entropic penalty for binding, an evolutionary adaptation that contributes to its high affinity for the receptor^6^. We further identified high probability molecular binding interactions that inform both vaccine design and therapeutic development, which may include recombinant ACE2-based spike decoys^7^ and/or allosteric S RBD-ACE2 binding inhibitors^8,9^ to prevent or arrest infection and thus disease.

## Infection of host cells by SARS-CoV-2 is initiated by binding of viral spike protein to angiotensin converting enzyme 2

The causative viral agent of the current worldwide COVID-19 pandemic, SARS-CoV-2 (Severe Acute Respiratory Syndrome Coronavirus 2), is transmitted by exposure of oral, nasal, and/or eye mucus membranes to SARS-CoV-2 viral particles. SARS-CoV-2, like SARS-CoV before it^10^, gains access to host cells by binding of its extensively glycosylated surface ‘Spike’ (S) protein receptor binding domain (RBD) to angiotensin-converting enzyme 2 (ACE2), a membrane-spanning protein that is expressed in epithelial cells of the gut, alveolar cells of the lungs^11^, as well as on other cell types, including macrophages^12^ of the host. A detailed description of S RBD-ACE2 interaction resulting in viral entry is provided by Wrapp *et al*. 2020, in their report on the cryo-electron microscopy (cryo-EM) structure of S pre-fusion^5^.

Spike is assembled as a trimer on the viral surface and it has been reported that at any given time, only one of the three S RBD monomers is accessible for ACE2 binding^13,14^ This monomer switching is a dynamic process that comprises hinge-like movements that either expose or hide the RBD^15–17^ Upon binding of the Spike S1 subunit to ACE2, the trimer is de-stabilized and the Spike S2 subunit is shed by cleavage predominately performed by the cell-surface serine protease TMPRSS2 or, to a lesser extent, pH-dependent endosomal proteases^3,18,19^ in a process referred to as priming. The S2 subunit then takes on a stable conformation post fusion^4^.

The endocytosis and fusion of the virus to the host cell results in release of the viral RNA sequences that then co-opt the cellular machinery to generate new viral proteins and synthesize viral RNA. This is followed by virion assembly and release by bursting and destruction of the host cell.

The S protein is not only required for viral entry to the host cells, but is also highly antigenic^4^ and thus considered a prime target for neutralizing antibodies, vaccines, and novel therapeutic treatments. For all of these approaches, it is essential to know as much as possible about interactions of the S RBD with ACE2 at the molecular level. The recently published crystal^20^ and cryo-EM^4,5^ structures of SARS-CoV-2 have been of great utility in this regard by providing atomic-scale resolution, but they do not reveal dynamic changes in protein interactions over time. This dynamic information can be attained by *in silico* Molecular Dynamics (MD) simulations and computational analysis.

Opportunities to find effective therapeutics or vaccines for COVID-19 can be greatly increased by MD simulation-enriched knowledge of predicted interactions between not only the 5 RBD and ACE2, but also between either of these proteins and potential vaccine-expressed antigens, antibodies, therapeutic proteins or chemical entities.

## Millisecond-scale Molecular Dynamic (MD) simulation provides accurate details of S RBD pre-fusion conformation and key residues for ACE2 interactions

Using the known sequence of S^21,22^ and recently reported cryo-EM structure of the S RBD-ACE2 complex^4,5^, we performed *in silico* MD simulation studies of free S RBD and the S RBD-ACE2 complex for 2.9 and 2 milliseconds (ms), respectively. The simulations utilized cutting-edge graphics processing units (GPUs) hardware in a large-scale cluster. The long timescales and overlapping behavior seen in the trajectories provides confidence in the biological relevance of the findings.

We used the PDB 6M17 S RBD-ACE2 complex structure as shown in Figure 1A for both free S RBD and S-RBD-ACE2 interaction MD simulations, wherein the residues used to calculate interface dynamics are highlighted and shown in an expanded view.

**Fig. 1.**
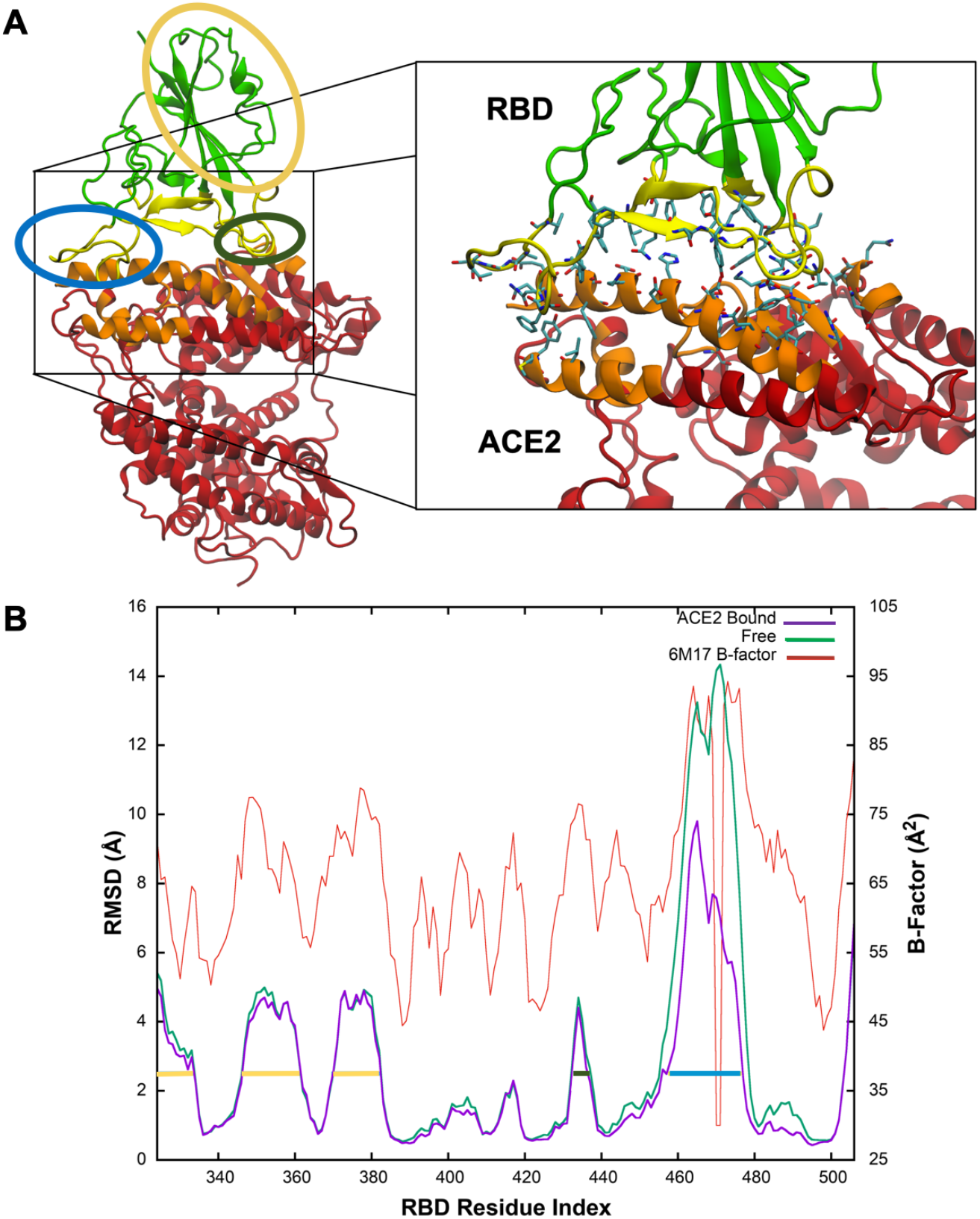
View of the ACE-2-RBD complex from PDB 6M17 and relative RMSD values. (A) The RBD-ACE2 complex structure (PDB 6M17) is shown with S RBD rendered in green and yellow while ACE2 is in red and orange. The zoomed view on the right shows residues found to frequently participate in intramolecular interactions. Regions circled in yellow and dark green show medium flexibility while those circled in blue show high flexibility. (B) Both the ACE2-bound (purple) and free RBD (green) simulations were aligned to the same structure so that RMSDs, calculated only for backbone atoms, are comparable. Regions with medium (yellow; residues ~336-345, 359-373, and 382-394 or dark green; residues 445-449) or high (blue; residues ~470-487) flexibility correlating with those shown in A are indicated by horizontal lines. B-factors (orange) from the original cryo-EM structure (PDB ID 6M17) are also shown on the second axis and indicate that the model follows the same trends as the experimental measurements.

To assess the relative conformational stability of free S RBD and the S RBD-ACE2 complex, we calculated the Root Mean Square Deviation (RMSD) for ACE2-bound RBD relative to the free RBD using only the backbone atoms, as shown in Figure 1B. RBD consists of a rigid core of residues (RMSD < 2.0Å) with flexible peripheral loops. A significant part of the binding interface is part of this rigid core, suggesting RBD has adapted to bind ACE2 with high affinity. The two interface regions that are not part of the core have medium and high RMSDs. Residues 445-449 form a loop whose moderate flexibility is not affected significantly by binding. In both the ACE2-bound and free RBD simulations, residues 470-487 comprise a highly flexible loop (yellow circle and bar in Fig. 1A and 1B, respectively). The dramatic shift in RMSD between bound and free RBD suggests that the loop contributes to the binding affinity. Finally, residues 336-345, 359-373 and 382-394 (blue circle and bars in Fig. 1A and 1B, respectively) are moderately flexible. While these residues are distant from the binding interface, they have been observed to form a binding pocket capable of accommodating linoleic acid^22^.

The MD simulations produced millions of conformations of the flexible complex making direct interpretation of the conformational landscape all but impossible. We used Principal Component Analysis (PCA) to condense most of the conformational variability into a few vectors. The resulting conformational probability densities for free (solvated) RBD (Fig. 2A-C) and ACE2-bound RBD (Fig. 2D-F) show good overlap (Fig. 2G-I), indicating that ACE2-bound RBD’s favored conformation overlaps well with one of free RBD’s major conformations. This suggests that free RBD’s flexible loop is conformationally primed to bind ACE2. Representative simulation structures from various regions of the probability distributions shown in Figure 2J. I, II and III are from the free (solvated) RBD simulation; Area I corresponds to the high-probability region that overlaps with the ACE2-bound maxima, II is from the most densely sampled region, and III was chosen because it represents a small density maximum in an otherwise sparsely populated region of conformation space. IV is from the ACE2-bound distribution and is similar to I. Both I and IV - each with an extended loop region - are high probability conformations for free or bound RBD.

**Fig. 2.**
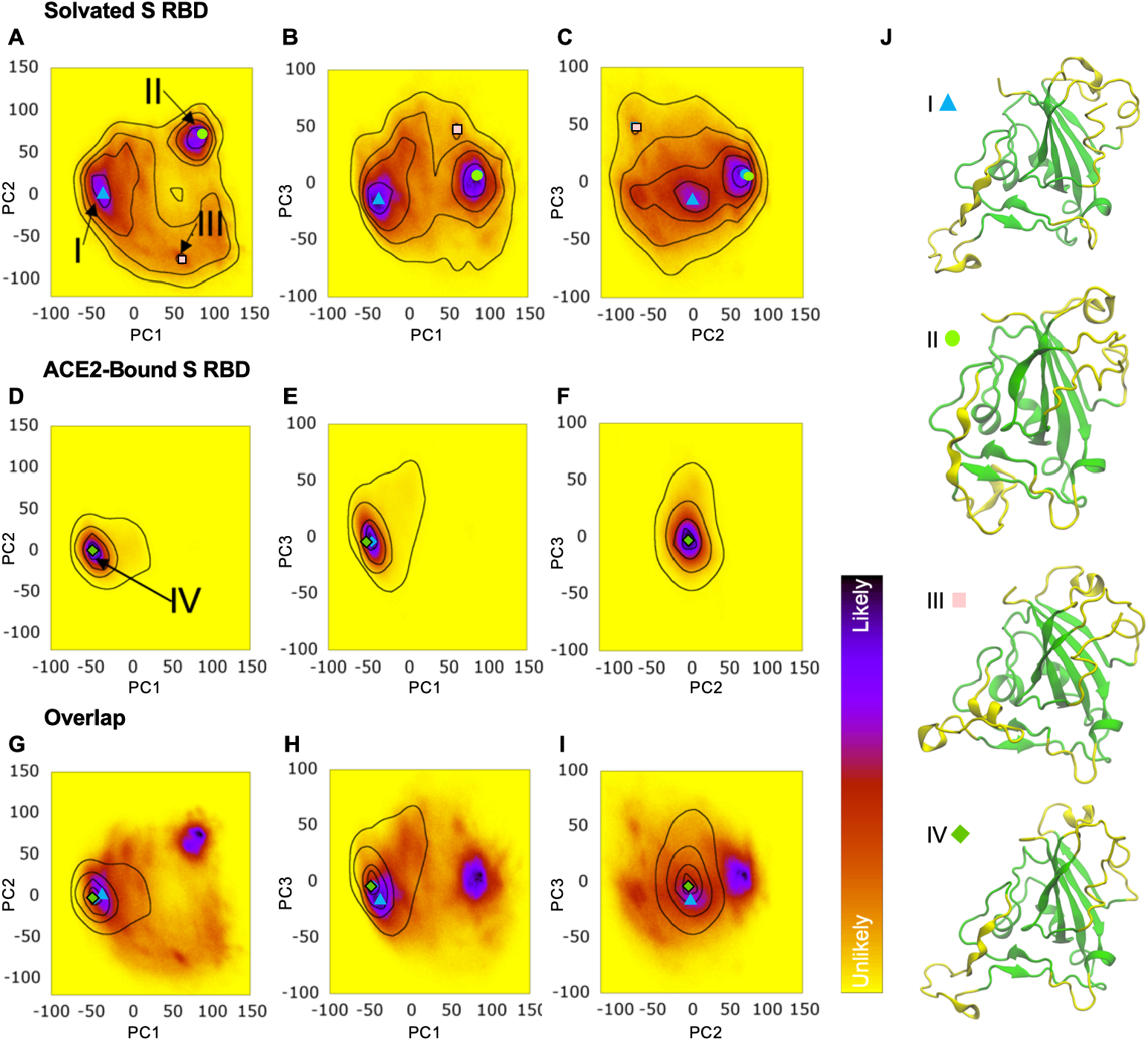
Principal component analysis (PCA) of RBD andACE2-SRBD dynamics. (A-I) Plots of the first three principal components calculated for the backbone of the whole RBD are shown. Eigenvectors were calculated from the combined set of ACE2-bound and free RBD conformations. PCA probability densities are shown for solvated (free) RBD (A-C), ACE2-bound RBD (D-F), and overlap (G-I), wherein the free RBD heatmaps and the ACE2-bound contours (outlines) are superimposed to show the overlap. (J) I - IV are representative simulation structures from various regions of the probability distributions; their locations are labeled in A and D and their marker shapes are represented in (A-I).

In addition to the PCA distribution calculations for S RBD in the free and bound forms described above, we calculated the PCA distribution for residues making up the interface between S RBD and ACE2. In Figure 3A, (PC1 and PC2), conformations I and II shown in Figure 3E are from the highest probability density; I is the cryo-EM structure of PDB 6M17, and II is representative of the majority of structures. The ACE-2 and S RBD binding surfaces have complimentary shape and charge distribution when bound. The simulations show that, when bound to ACE2, RBD exists almost entirely near a single conformation. However, on the millisecond timescale, we do see the flexible loop sampling a range of conformations. To convey the extent of conformation space explored, the raw PCA data for the interface interactions is shown in Supplementary Figure S1.

**Fig. 3.**
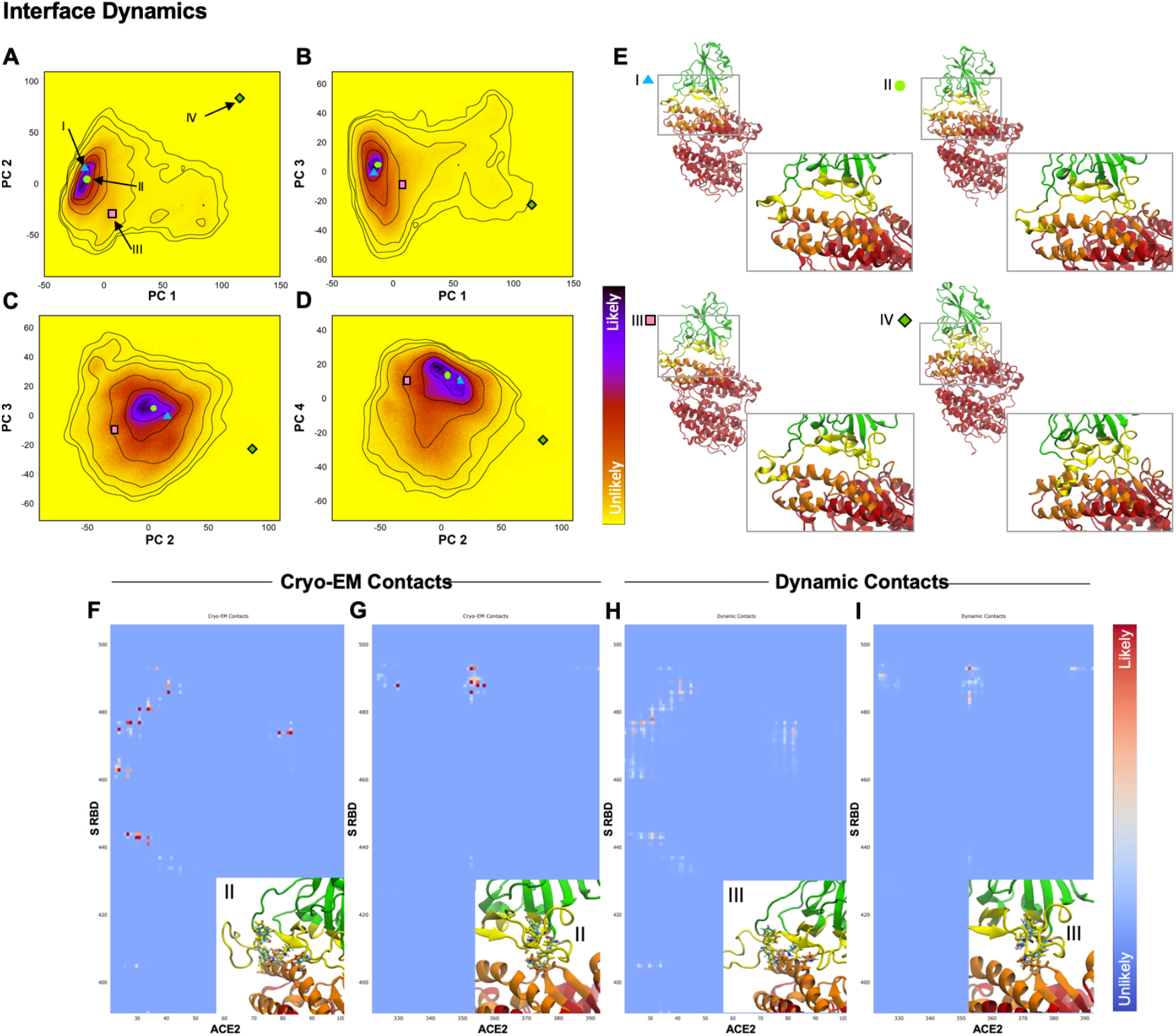
Intramolecular dynamics and contact strength between SRBD and ACE2 predicted by cryo-EM versus MD simulation. Heat maps of PCA distributions calculated for residues making up the interface between RBD and ACE2 are shown for (A) PC1 and PC2, (B) PC1 and PC3, (C) PC2 and PC3, and (D) PC3 and PC4. (E) Structures indicated in (A) are shown with a zoomed interface (box). For PCA, darker colors indicate higher probability. Iso-probability contours were calculated by smoothing the probability data using a distance-weighted grid averaging. (F-I) The strengths of residue interactions based on cryo-EM for structure II (inset) and for MD simulation for structure III (inset) are represented. Color scale is consistent across images, with darker red indicating greater strength of interaction. Backbone ribbons are colored as in Fig. 1A. Several strongly contacting residues are rendered as sticks. The residues colored by atom type are the cryo-EM coordinates. The orange and yellow residues are the same residues but from the simulation.

The timely determination of the spike RBD cryo-EM structure by Walls *et al*.^4^ and Wrapp *et al*.^5^, not only confirmed that the RBD of the spike protein of SARS-CoV-2, like SARS-CoV, bound to ACE2 but did so with higher affinity^5^ and revealed the presence of a furin cleavage site between the S1-S2 subunits that distinguished it from SARS-CoV^4^ Further, Wrapp’s 3.5-angstrom resolution of the cryo-EM structure of the S trimer in the prefusion conformation also showed that in the predominant state, only one of the three binding domains is rotated up in the accessible position. Their cryo-EM structure very likely contributed to the design of vaccines under development. In our efforts here, we hoped to build on the initial cryo-EM analyses with a closer look at the RBD-ACE2 interface using MD simulation. Not unexpectedly, by using molecular dynamics, we found a much broader range of interactions than the static cryo-EM structure would suggest. Figures 3F – I show the measured likelihood of interface contacts in the cryo-EM structure (Fig. 3F and G) and the 2 ms MD simulation (Fig. 3H and I). To compare our MD simulation findings for interface interactions to cryo-EM-based analyses, we used the full set of equilibrium simulations to determine the strength of intramolecular contacts and found that cryo-EM contacts are over-estimated (Fig. 3F and G) as reflected by the darker red points in the plot compared to MD simulation (Fig. 3H and I). MD simulation at the duration used here reveals many contacts that while weaker, are still significant. Thus, the simulation provides a more complete and accurate picture of key interactions. The strongest contacts revealed by cryo-EM were from structure II shown in Figure 3E, whereas those seen by MD stimulation were from structure III. While the residues interacting are similar for cryo-EM and MD simulation, the specific conformations (II versus III, respectively) during the highest affinity binding are different.

To better understand the rare events involved in the binding process, we performed Adaptively Biased MD (ABMD) simulations^23^. ABMD applies a force to encourage the system to sample new conformations. The biasing force is applied to collective variables (CV) that are themselves functions of the atomic coordinates. The CVs used here were the number of intramolecular contacts and the distance between the centers of mass (COM) of the S RBD and ACE2 interface regions. The resulting free energy surface is shown in Figure 4A. Structures corresponding to points in the most likely (i.e. lowest free energy) path through the free energy surface (Fig. 4A) are shown in Figure 4B. The S RBD interface surface comprises the region extending from the flexible loop region indicated by the blue arrow in Fig. 4B, structure II to a second, relatively extended unstructured loop region that forms a hydrogen bond with ACE2 residues at a hydrophobic interface, indicated by a black arrow in the same structure. It is this latter unstructured loop region that binds most tightly and is the last region to dissociate. This relatively high affinity is due to several hydrogen bonding and hydrophobic interactions. For example, Lys353 in ACE2 participates in four of the twelve most frequently observed H-bonds and three of the eight most frequent non-polar contacts. The most frequent H-bond is between the carbonyl oxygen of Lys353 and the backbone amide of Gly502 from RBD, while the most frequent non-polar interaction is between the aliphatic carbons of Lys353 and the aromatic ring of RBD Tyr505. The residues (ACE2 326 and 353-357, RBD 445-449, 502 and 505) involved in this interaction have been experimentally shown to be necessary for binding^24^.

**Fig. 4.**
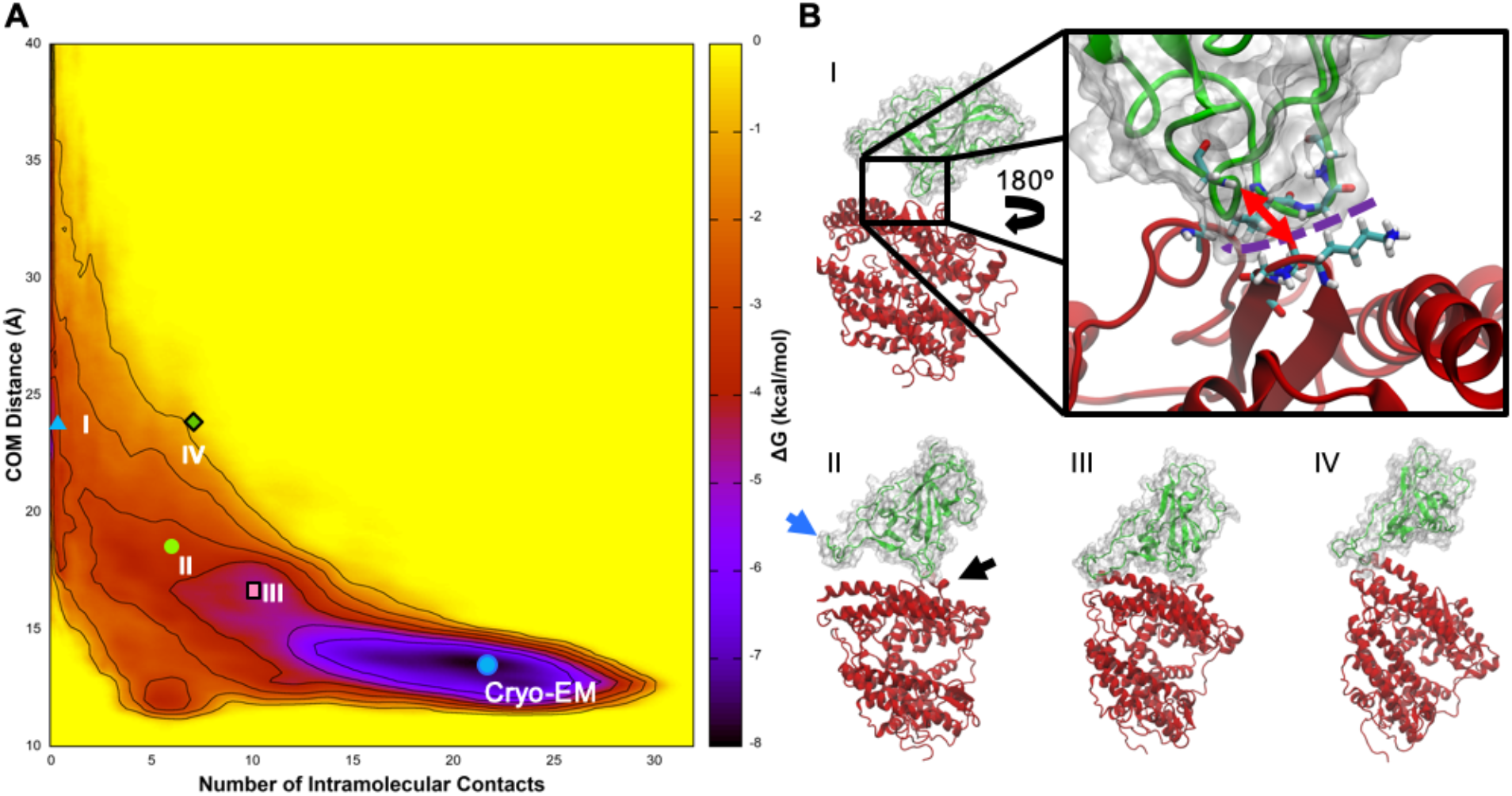
Free energy of dissociation of S RBD and ACE2. (A) The blue dot is the cryo-EM structure (PDB ID 6M17). Points I-IV correspond to the structures in (B). I-III are representative of the most likely path through the free energy surface. The inset of I shows the hydrogen bond (red arrow) and hydrophobic surface (dashed purple line) responsible for the high affinity unstructured loop affinity. Note that the inset has been rotated around the z-axis by ~180 degrees compared to the zoomed out view.

An example from the ABMD simulation of a typical binding event along the main pathway can be viewed in the video provided in *Supplementary Materials*. In particular, the induced fit of the flexible loop is apparent, whereas the high-affinity unstructured loop region is relatively rigid in the binding-ready conformation.

Structure IV in Figure 4 represents a minor dissociation path that lies in the higher-energy region above the main pathway defined by I, II, and III in Figure 4A. These structures are typical of conformations in this part of the free energy surface. In IV, the high-affinity unstructured loop is dissociated while the flexible loop remains bound. This represents an alternative pathway for association where the flexible loop makes contact first. While paths through this region are less probable, they are still relevant to binding and dissociation.

To ensure that the applied biases did not distort the interface structure, we compared the interface PCA distributions for the ABMD and equilibrium simulations as shown in Supplementary Figure S2. This shows that the ABMD simulations are sampling similar conformational space compared to the equilibrium ACE2-bound RBD simulations. This control assures the RBD structure has not been distorted by the biasing forces acting upon it during ABMD.

## DISCUSSION

### The free spike receptor binding domain adopts an optimized, ACE2 binding-ready conformation facilitating binding and infection

While others have performed MD simulations for S RBD-ACE2 or peptides that interact with spike or ACE2 to prevent binding,^25–28^ to our knowledge, MD simulations of both free and bound S RBD for the durations have not been previously reported. These millisecond scale MD simulations provide remarkably strong support for the findings described herein.

Perhaps the most important finding from an evolutionary standpoint is that free S RBD adopts an optimized, ACE2 binding-ready conformation, incurring little entropic penalty for binding with ACE2. This is in accord with and provides an explanation for the experimentally observed (and herein confirmed and detailed) high affinity intermolecular interactions between spike RBD and human ACE2^6^. This binding-ready RBD very likely contributes significantly to the virus’ high infectivity, particularly when taken into consideration with other factors that contribute to its rapid spread such as the mode of transmission (airborne and through surface contamination) and stability of the virus on surfaces/in air. Finally, the ACE2 presents a target that is abundant both in the respiratory and digestive tracts and thus readily available to the virus.

As mutants of the virus inevitably appear^29^, particularly those with variant spike sequences^30^ such as the Y453F variant recently found in mink^31^, application of the deep MD simulation could help predict infectivity and evolutionary forces (such as variants of ACE2 in certain populations) that may affect emergence of these mutants.

### Application of millisecond-scale MD simulation to design of vaccines and therapeutics

Of immediate utility in addressing the pandemic, the high probability molecular binding interactions identified here can be considered in both vaccine design and therapeutic development.

The presented MD simulation-derived information has the potential to expand on the insights gained from the cryo-EM structure and thereby contribute to vaccine design. Antibodies against the high-affinity S RBD-ACE2 interaction sites described here are likely to be highly neutralizing and therefore relevant to the screening and design of recombinant neutralizing antibodies (Nabs). These residues and structures are also likely to be evolutionarily conserved, decreasing the likelihood of escape mutations. The specific use of the MD simulations should be extremely valuable for *in silico* design and screening of potential Nabs.

Another therapeutic approach to prevention of SARS-CoV-2 infection that would likely benefit by consideration of these high-affinity residues is the ‘spike trap’ (or decoy) approach^7,32^. Recombinant ACE2 sequences could potentially compete with endogenous host ACE2 binding of S RBD, thus preventing infection. The optimal trap would not just comprise a sequence with the highest affinity for S RBD, but one with a low affinity for natural host ligands of ACE2, such as angiotensins 1 and 8. This would produce a selective trap that reduces the rate of the infection and doesn’t interfere with normal host functions. The analyses performed here could be adapted to identify sequences with these characteristics.

Long-duration modeling can also be applied to *in silico* screening of molecules that could inhibit the S RBD-ACE2 interaction^8,9,33^. These may be antibodies, peptides or small molecules that either bind directly to the interaction surface or act allosterically to stabilize S RBD in a conformation with low ACE2 binding affinity.

Regardless of the approach to development of therapeutics to address the COVID-19 pandemic, the detailed information reported here with high confidence due to the extraordinary duration of MD simulation of both free and ACE2-bound RBD provides an opportunity to effectively expand our arsenal against this disease, provides a method for identification of spike mutants that pose possible new threats, and highlights the feasibility and merits of what previously might have been considered an ambitious undertaking.

## Supporting information

RBD-ACE2 interaction video

## ACKNOWLEDGEMENTS

We would like to thank the team at Microsoft Corporation including the teams at Azure and Microsoft Research. Jim Jernigan, Peter Lee, James N. Weinstein, Kris Zentner, Benjamin Huntley, Ian Finder, Jack Kabat, Katie Weissenfels, Jordan Boland, Desney Tan, and Oleg Losinets allowed us to avail ourselves of their computing power and provided invaluable technical support.

## SUPPLEMENTARY MATERIALS

### Methods

#### System Setup

The WT-ACE2/RBD complex was built from the cryo-EM structure, PDB 6M17 of fulllength human ACE2 in the presence of the neutral amino acid transported B^0^AT1 with the S RBD as shown in Yan *et al*.^21^ using RBD residues 336-518 and ACE2 residues 21-614. Protein models came from the Amber ff14SB force field^34^. Solvating waters were packed around the complex using the RISM program from AmberTools19^35^. This structure was then solvated with 44,754 additional TIP3P water molecules and neutralized with 23 sodium ions. To build the system, we used tleap from AmberTools19. Molecular dynamics (MD) simulations were run with the GPU-enabled version of Amber18, pmemd.cuda^35^. Multiple copies of the initial starting structure were minimized and equilibrated with unique random seeds. After minimization and heating to 300K, system pressure was relaxed to 1 atm for 20 ns in the NPT ensemble.

As a comparison to its dynamics in the complex, unbound RBD was also built and simulated. The initial structure was also residues 336-518 from PDB 6M17. RISM was used to place water molecules in favorable contact with RBD and then 14024 additional waters were added. Two chloride ions neutralized the system. Equilibration followed the same protocol as the complex.

#### Simulation

In order to both explore and adequately sample conformation space, a two phase procedure was used. In the first phase, temperature replica-exchange molecular dynamics (tREMD)^36,37^ was used to encourage the complex to explore new conformations. The second phase used equilibrium molecular dynamics simulations to thoroughly sample the conformation space explored by tREMD. All simulations were run with a 2 fs timestep and an 8Å cutoff. All post-equilibration simulations were run in the NVT ensemble.

For each equilibrated structure, 40 tREMD replicas were used. Replicas were spaced in 1 degree increments from 300 to 339K and swaps were attempted every 10 ps. Simulations were performed with the MPI version of GPU-enabled pmemd, pmemd.cuda.MPI. After 1 ms aggregate tREMD simulation time, frames at 300K were extracted from the full simulation trajectories. These trajectories were processed with PCA and clustered using the agglomerative algorithm with the k-means metric in cpptraj from AmberTools19. Representative frames from the resulting clusters were used to start simulations for the second phase of sampling.

Equilibrium MD was run with pmemd.cuda and produced a total of 2 ms of sampling at 300K. The resulting trajectories were analyzed to provide the equilibrium results presented here.

Simulation of the free RBD molecule followed the same protocol as the complex. A total of 261.6 μs of data was collected for tREMD and 2.9 ms for equilibrium.

#### Per-Residue Root-Mean-Square-Deviation (RMSD)

To compare the flexibility of bound vs. free RBD, equilibrium trajectories for both systems were first stripped of everything other than RBD. The cryo-EM structure was then used as a reference to which the backbones of both the bound and free simulation structures were aligned. RMSD was calculated from the total dataset for each residue without realigning. This first pass gave an indication of which residues were flexible and which were not. To get a clearer picture of how per-residue flexibility is impacted by ACE2 binding, residues with an RMSD less than 2.5Å in the first pass were defined to be structural. In the second pass, the backbones of only the structural residues were aligned to the cryo-EM structure and then the RMSD was again calculated for all RBD residues without realigning. In the second pass, the bound and free datasets were analyzed independently so that the differences could be seen. The plot was generated using gnuplot.

#### Principal Component Analysis (PCA)

Both the RBD comparison and RBD/ACE2 interface PCA plots were created with the following protocol. The only differences were the input data and the atom selection. The RBD comparison used RBD from both the ACE2-bound and free datasets and was calculated from the backbone atoms of all residues. The interface analysis used only the RBD/ACE2 complex data and used the interfaces as shown in the text. PCA plots were calculated using cpptraj.

Eigenvectors were calculated by first aligning simulation structures to the cryo-EM structure. The same set of atoms was used for both the alignment and the PCA calculation. PCA projections were calculated for bound and for free RBD separately. PCA distribution plots were generated with gnuplot. Iso-probability contours were generated by smoothing the input data on a distance-weighted grid.

#### Contact Analysis

Intermolecular contacts were measured using the ‘nativecontacts’ functionality in cpptraj. The intermolecular region defined for PCA was used to search for contacts. The contact map was generated using the “byresidue” option to generate output on a per-residue basis. The raw output was processed using command-line tools and plotted in gnuplot.

#### Potential of Mean Force (PMF)

Based on exploratory simulations, the center of mass (COM) distance and number of intermolecular contacts were chosen as the two collective variables (CVs) best suited to overcome un/binding barriers. The two centers of mass were defined as the interfacial alpha carbons from either RBD or ACE2. The “contacting residues” were all intermolecular pairs from the contact analysis for whom the sum of the contact fraction was greater than 50.

Initial structures for the 2D PMF were calculated by running 1D steered MD (SMD) from 10 independent equilibrium structures. SMD was run only on the contact number CV. From the initial CV value, the number of contacts was smoothly pulled to 0 over a 10 ns simulation. Starting structures were randomly selected from these trajectories.

The 2D PMF was calculated using well-tempered multiple-walker adaptively biased MD (ABMD) as implemented in Amber18. The timescale was set to 100.0 ps and the well-tempered temperature was set to 1500.0 K. ABMD simulations were run for a total of 32 μs. The PMF was plotted with gnuplot. PCA plots for the interface during dissociation were calculated from the eigenvectors generated from the equilibrium interface PCA calculations.

### Supplementary Video

The video file of S RBD-ACE2 interaction, render_RBD-ACE2_binding_WW_opacity-0.3.mp4, is uploaded as Supplementary material.

### Supplementary Figures

**Fig. S1.**
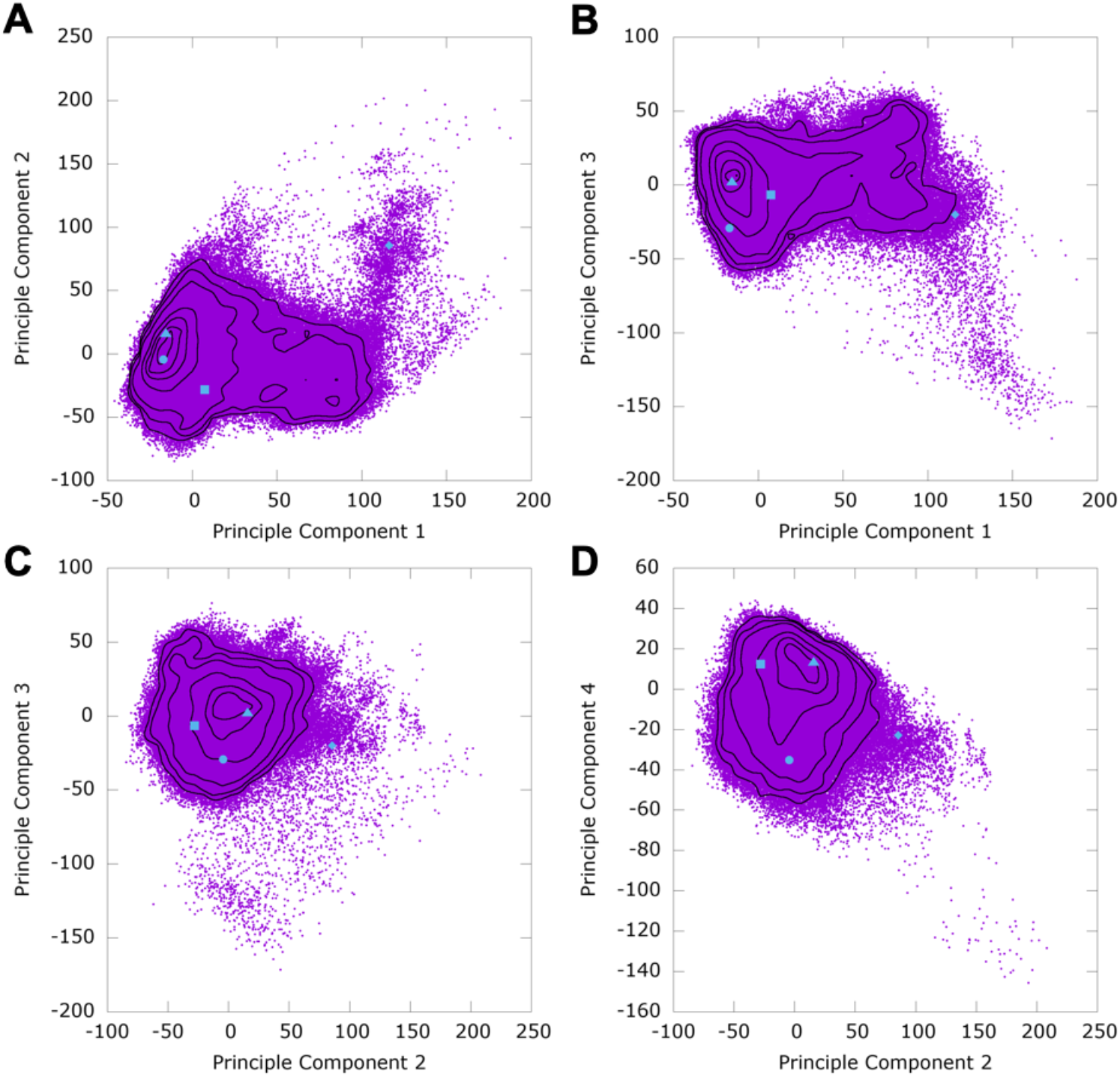
Raw PCA output. The raw PCA output is shown for interface interactions from main Figure 2 K-N to reveal the full extent of sampling. The same contours from the heatmaps are shown for comparison; here A = Fig. 2K, B = Fig. 2L, C = Fig. 2M, and D = Fig 2N.

**Fig. S2.**
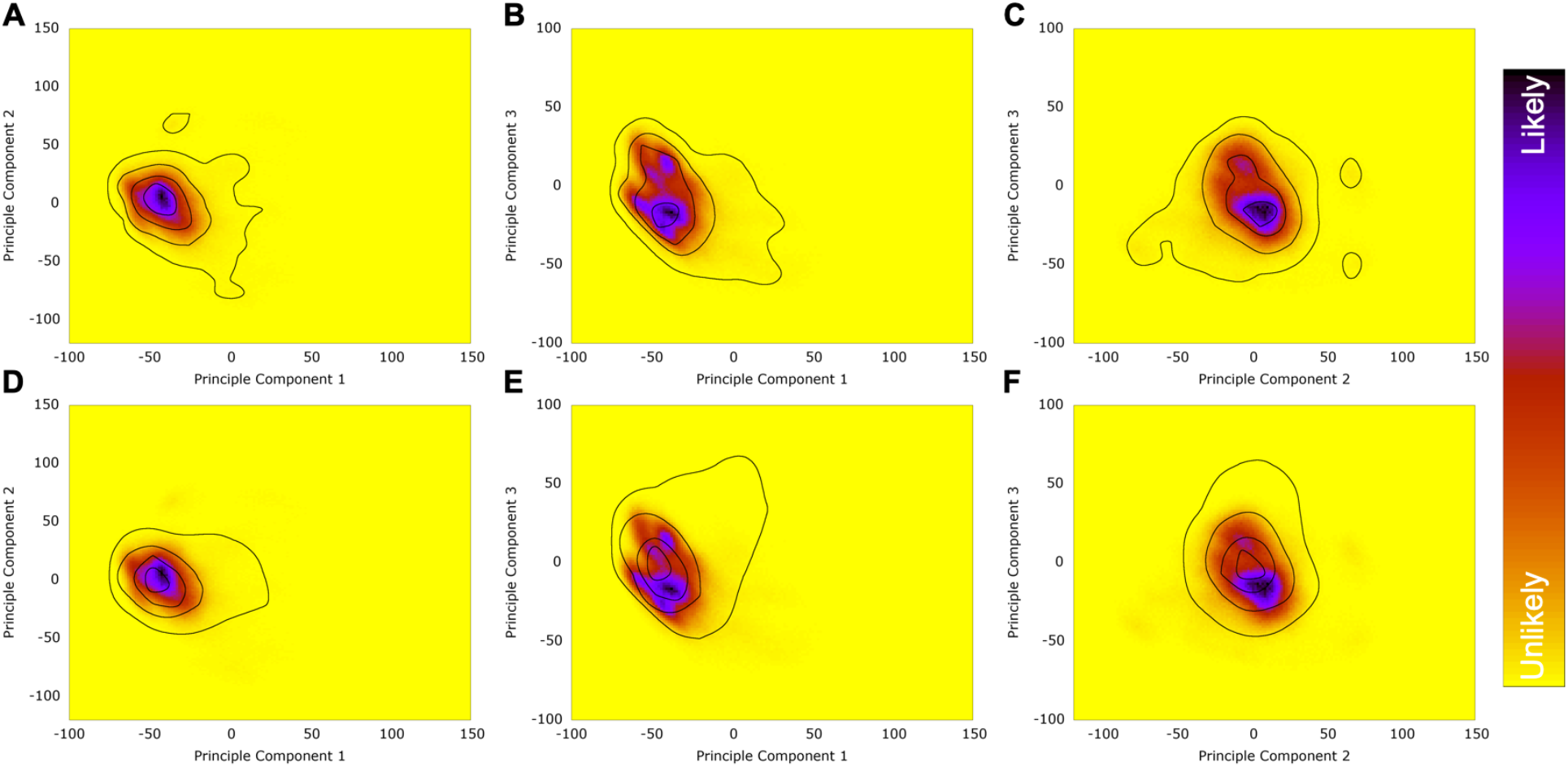
Distributions calculated by projecting the adaptively biased molecular dynamics (ABMD) RBD structures onto the PCA space for ACE2-bound RBD simulations. (A-C) Plots of contours for the ABMD simulations. (D-F) Plots of contours calculated from the 2 ms equilibrium data for ACE2-bound RBD simulations superimposed on the ABMD distribution heatmaps. This shows that the ABMD simulations are sampling similar conformation space compared to the equilibrium ACE2-bound RBD simulations. ABMD simulations of the complex binding and unbinding did not significantly distort RBD compared to free RBD.

## Notes

### Competing Interest Statement

The authors have declared no competing interest.

